# Activation of mitochondria is an acute Akt-dependent response during osteogenic differentiation

**DOI:** 10.1101/2020.06.22.164723

**Authors:** C. Owen Smith, Roman Eliseev

**Affiliations:** Center for Musculoskeletal Research, University of Rochester School of Medicine & Dentistry

**Author notes:** Corresponding author: Roman A. Eliseev, Center for Musculoskeletal Research, University of Rochester, Rochester NY 14624.

**Keywords:** Mitochondria, Osteoblast, differentiation, Akt, Wnt, BMP2, bioenergetics, osteogenisis

## Abstract

Osteogenic differentiation, the process by which bone marrow mesenchymal stem/stromal (a.k.a. skeletal stem) cells and osteoprogenitors form osteoblasts, is a critical event for bone formation during development, fracture repair, and tissue maintenance. Extra- and intracellular signaling pathways triggering osteogenic differentiation are relatively well known; however, the ensuing change in cell energy metabolism is less clearly defined. Here we tested the effect of osteogenic media containing ascorbate and β-glycerol phosphate, or various osteogenic hormones and growth factors on energy metabolism in long bone (ST2)- and calvarial bone (MC3T3-E1)-derived osteoprogenitors. We show that osteogenic media, and differentiation factors, Wnt3a and BMP2, stimulate mitochondrial oxidative phosphorylation (OxPhos) with little effect on glycolysis. The activation of OxPhos occurs acutely, suggesting a metabolic signaling change rather than protein expression change. To this end, we found that the observed mitochondrial activation is Akt-dependent. Akt is activated by osteogenic media, Wnt3a, and BMP2, leading to increased phosphorylation of various mitochondrial Akt targets, a phenomenon known to stimulate OxPhos. In sum, our data provide comprehensive analysis of cellular bioenergetics during osteoinduction in cells of two different origins (mesenchyme vs neural crest) and identify Wnt3a and BMP2 as physiological stimulators of mitochondrial respiration via Akt activation.

## Introduction

Mitochondria are dynamic organelles central to cellular energy production, metabolite synthesis, ion homeostasis, reactive oxygen species (ROS) generation and scavenging, epigenetic signaling, and apoptotic signaling in nearly every cell type (McBride, Neuspiel and Wasiak, 2006; Nunnari and Suomalainen, 2012; Schorr and van der Laan, 2018). The current understanding of stem cell energetics holds that following differentiation, cell reliance on oxidative phosphorylation (OxPhos) increases over glycolysis. Indeed, increasing the activity of glycolytic pathways is a common mechanism to induce stemness in cells *in vivo* and in induced pluripotent stem cells that revert back to glycolysis to become more stem-like. Therefore the importance of mitochondria in cell types classically considered to be glycolytic such as those of stem cell niches is a nascent field of research (4). The most obvious hypothesis for this metabolic transition is the approximately 16-fold increased energetic efficiency of OxPhos compared to glycolysis in ATP synthesis allowing for heightened rates of cellular activity to meet the demands of differentiation. Alternatively, mitochondria are central hubs of cellular second messengers and epigenetic signals (5–7). Bone marrow mesenchymal stem/stromal cells (BMSCs, a.k.a. skeletal stem cells) are somatic multipotent progenitors capable of differentiation into bone-forming osteoblasts (OBs), cartilage-forming chondrocytes, and fat-forming marrow adipocytes (8). The heightened energy demand from the osteoblast, compared to the other lineages, is due to high levels of collagen, bone matrix protein, biosynthesis followed by mineral deposition (9). Recent work by our group and others demonstrate that OxPhos is important in osteogenically differentiating BMSCs and their progeny, OB and osteocytes, as measured by an increase in oxygen consumption (10–12). It has been proposed that rather than increased OxPhos, the increase in oxygen consumption is a result of an increased demand for NADH in order to maintain the activity of lactate dehydrogenase as a result of increased glycolysis, an increase in ribose precursor production, or an increased demand for ATP. Currently, it is unclear what role metabolic plasticity plays in OB differentiation and development. Increases in OxPhos in BMSCs (10,12,13) and calvarial osteoblasts (11) have been reported; however, there remains the unanswered possibility that the observed increase in oxygen consumption is a result of decreased coupling efficiency or increases in non-mitochondrial oxygen consumption. There is also the possibility that long bone and calvarial bone OB metabolism are biologically distinct (14). Conversely, there are reports that OxPhos is not activated during osteogenesis in long bone-derived cell lines (15), calvarial-derived cell lines (16), and primary BMSCs (17), however, these studies describe changes in oxygen consumption under experimental conditions which contain pyruvate, supraphysiological levels of glucose, or both. As pyruvate can feed directly into the mitochondrial citric acid cycle and high glucose is a powerful driver of the Crabtree effect, these conditions could bypass any constraints that specifically affect OxPhos. The metabolic plasticity of mitochondria allows for the heterogeneous mix of substrates classically used in cell media and mitochondrial buffers: glucose, pyruvate, and glutamine to substitute as carbon entry points in the Krebs’ cycle (11,18). Glutamine is converted to α-ketoglutarate to enter the Krebs cycle through the process of glutaminolysis and is used by OBs as an alternate fuel source (19). Lastly, bioenergetic characterization of the specific cause of cellular changes in oxygen consumption by mitochondrial versus non-mitochondrial respiration, respiration directed toward ATP synthesis, and uncoupled respiration (20) is often overlooked in these assays. We have previously demonstrated that under the most physiologically relevant cell culture conditions, OxPhos is indeed activated in BMSCs incubated in osteogenic media composed of 50 μg/mL ascorbate and 2.5 mM β-glycerol phosphate in the presence of low (i.e. physiological 5 mM) glucose (12). We found that this process is not accompanied by major gene expression or mitochondrial mass changes but is accompanied by changes in mitochondrial dynamics. Concurrent work by the Kowaltowski group (21) reported similar results. The trigger(s) and upstream signaling for the observed mitochondrial changes during osteoinduction remain unclear and are the focus of this work.

In this study, we comprehensively explore the link between mitochondrial activity and osteogenic differentiation of two distinct long bone- or calvarial-derived osteoprognitor cell lines (ST2 and MC3T3-E1, respectively), applying either commonly used but artificial osteogenic media components or various physiologically relevant osteoinduction factors. Our data demonstrate that acute stimulation of mitochondrial oxygen consumption by osteogenic media, Wnt3a, or BMP2 upon osteoinduction is a common feature in both long bone- and calvarial-derived osteoprogenitors. We further show that this mitochondrial activation is completely dependent on activation of Akt. Intriguingly we observe that both the time course and degree of change in mitochondrial parameters are distinct depending on the osteogenic origin and mechanism of osteoinduction.

## Results

### Long bone-derived ST2 and calvarial bone-derived MC3T3-E1 osteoprogenitors require mitochondrial activation in osteogenic media

ST2 and MC3T3-E1 are mouse derived osteoprogenitor cell lines that do not have high alkaline phosphatase (ALP) or mineralization activities in non-differentiating growth media but become strongly positive for ALP after 2 days and for mineralization after 7 days of incubation in osteogenic media (22,23). These changes are concomitant with gene expression for early and late markers of osteogenic differentiation as measured by real-time RT-PCR (Figure 1A, B). As changes in mitochondrial activity are gaining attention as a hallmark for cellular reprogramming (24); we first asked whether osteogenic differentiation required mitochondrial activity. ST2 and MC3T3-E1 cells were incubated with low doses (1/10^th^ required for complete inhibition) of various metabolic inhibitors (0.1 µg/ml Oligomycin (Oligo), 0.2 µM (FCCP), 0.1 µM Rotenone (ROT), or 0.1 µM Antimycin A (AA)) for 48 hours in non-differentiating growth media, then induced to undergo osteogenesis by addition of ascorbate and β-glycerolphosphate (*i*.*e*. Osteoblast Media: OBM). At these concentrations, none of the inhibitors impacted cell viability as assayed by crystal violet (CV) stain in the undifferentiated state (Figure 1C, D “day 0”). Following addition of ascorbate and β-glycerol phosphate (OBM), both cell types proved highly sensitive to addition of mitochondrial ATP synthesis (Oligomycin) or electron transport chain inhibitors (Rotenone, Antimycin A) but not to mild uncoupling (FCCP, Figure 1C, D “day 4”). This demonstrates the importance for mitochondrial respiratory activity on osteoprogenitors at the point of initiating the osteogenic program.

**Figure 1:**
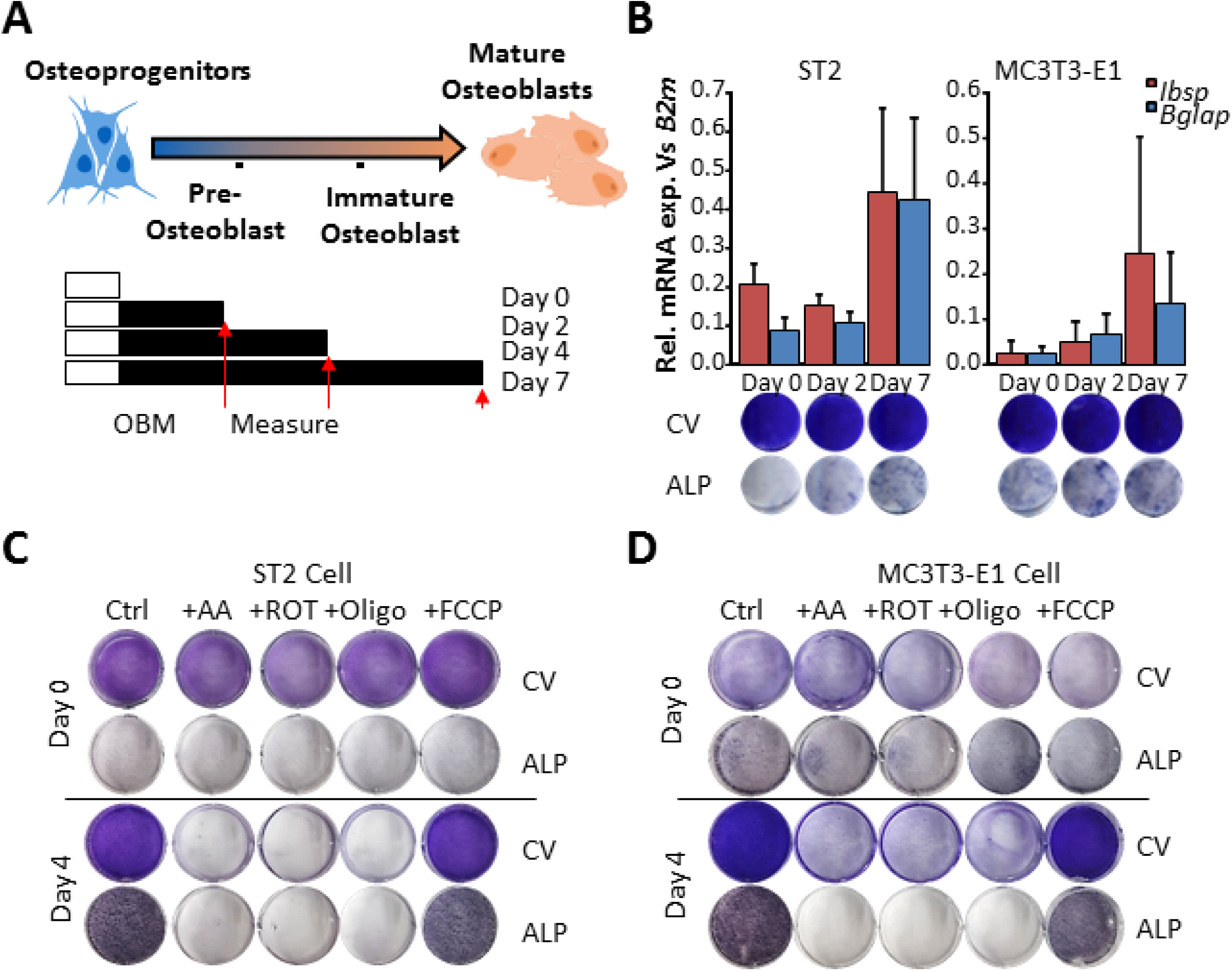
cell differentiation in osteogenic media activates mitochondria. (A) Top: Schematic representation of osteoprogenitor cell differentiation to OBs. ST2 and MC3T3-E1 cells represent osteoprogenitor stage cells of the long bone and calvarial bone, respectively (cell images generated using biorender.com). Bottom: Schematic of time course. Cells grow to confluency (white bars) then are treated with osteoblast differentiation media (OBM, black bars) for 2, 4 or 7 days before measurements are taken (red arrows). (B) Expression of mRNA levels of OB markers *Ibsp* and *Bglap* relative to control *B2m* increases with differentiation. Control Crystal Violet (CV) staining and OB marker alkaline phosphatase (ALP) for same time points showed below. Metabolic inhibitors were added 48 hours prior to “day 0” time point (Antimycin A: AA, rotenone: ROT, Oligomycin: Oligo, carbonyl cyanide p-trifluoromethoxyphenylhydrazone: FCCP). ST2 (C) and MC3T3-E1 (D) cells stained at Day 0 and Day 4 following OBM induction and stained for CV or ALP. Data are means ± SD (n=5 for qPCR). Staining images are representatives of 3 replicates.

### Osteogenic induction acutely activates mitochondrial bioenergetic parameters

Given that cells showed sensitivity to mitochondrial perturbation only upon osteoinduction, we asked whether mitochondrial activation was an early event during osteogenic differentiation. Moreover, previously published studies analyzed the effect of osteogenic differentiation on mitochondria only at conventional time points post osteoinduction (i.e. 4, 7, 14 days); thus, the immediate bioenergetic responses remain unclear. We therefore used the XF96 Seahorse Flux Bioanalyzer to measure ST2 and MC3T3-E1 cell oxygen consumption rates (OCR) and extracellular acidification rates (ECAR) under standard and osteogenic conditions. This method allows for real time monitoring of cellular OCR as a secondary measure of mitochondrial respiration and ECAR as a secondary measure of glycolysis. The serial incorporation of specific metabolic inhibitors to block mitochondrial ATP production (Oligo), uncouple mitochondrial respiration (FCCP), inhibit the election transport chain (Antimycin A+Rotenone, AA/Rot), or to inhibit Glycolysis (hexokinase competitive inhibitor 2-Deoxyglucose, 2DG) allow for the calculation of aerobic and anaerobic contributions to cellular metabolism (see methods, Figure 2A, B). As previously reported for BMSCs, undifferentiated ST2 and MC3T3-E1 osteoprogenitor cell lines relied marginally on oxidative phosphorylation, showing a slight preference for anaerobic metabolism (Figure 2C, D white shapes). However cellular respiration was significantly and acutely increased in cells upon addition of osteogenic media; and as little as 1 hour was sufficient to maximally stimulate mitochondrial OCR in ST2 cells (Figure 2C), while MC3T3-E1 cells showed progressive increase in OCR up to 12 hours (Figure 2D). Measuring the slope of a line drawn between the basal and maximal rates shows a slope greater than 1 for all osteoinduced cells, indicating a bias towards aerobic respiration as the main driver of the heightened energetic state (Figure 2C, D inset).

**Figure 2:**
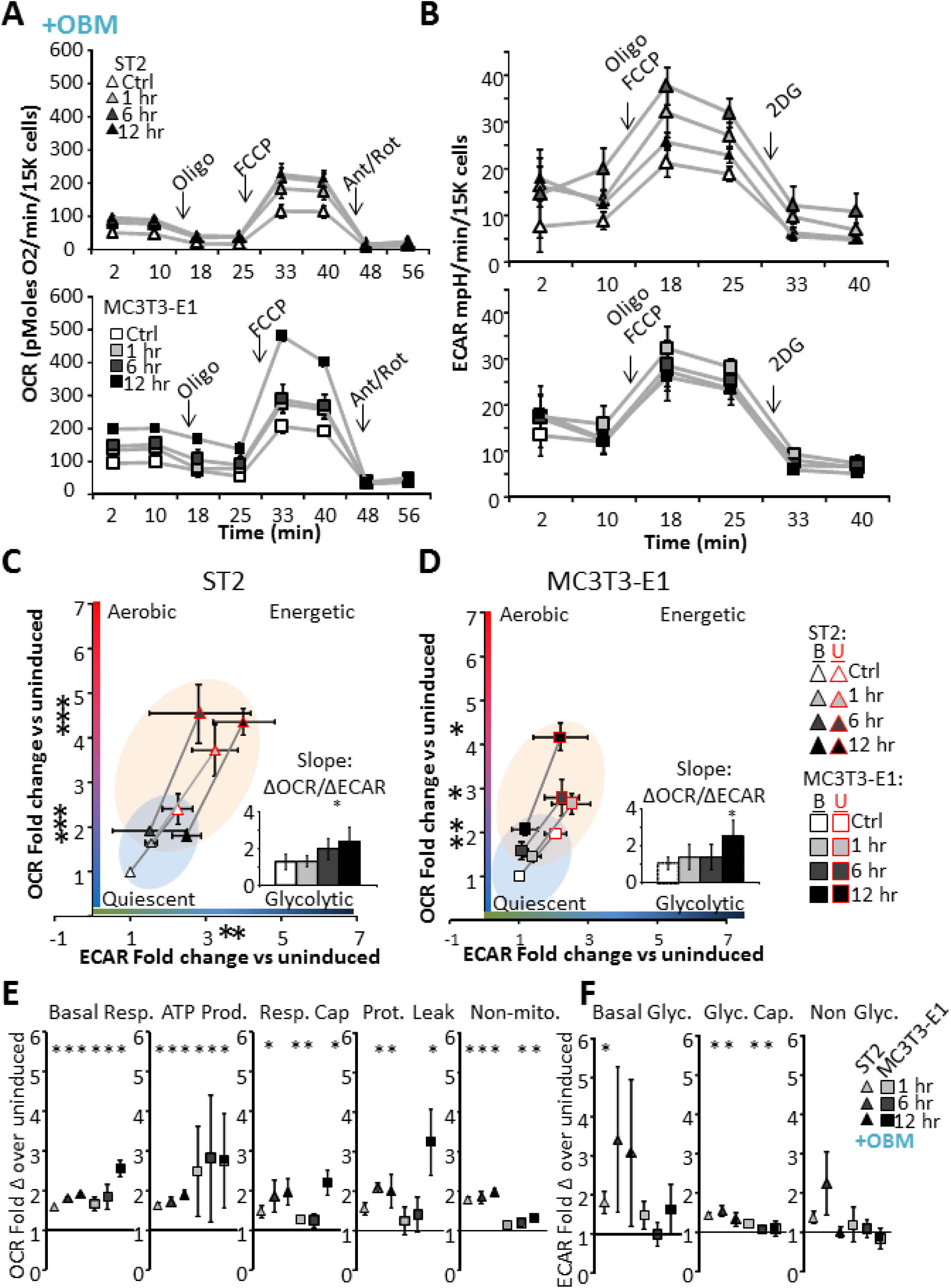
Bioenergetics in osteoprogenitor ST2 and MC3T3-E1 cell lines are acutely responsive to osteoinduction. (A) Oxygen consumption rates (OCR) is measured as an endpoint measure of mitochondrial respiration shown here for cells as an acute response to OBM for ST2 (top) and MC3T3-E1 (bottom) cells. Arrows indicate injection of metabolic modifiers to inhibit mitochondrial ATP synthesis (Oligo), uncouple mitochondria (FCCP), and inhibit electron transport chain (AA/Rot). (B) Extracellular acidification rates (ECAR) is measured as an endpoint for anaerobic glycolysis shown here for cells as an acute response to OBM for ST2 (top) and MC3T3-E1 (bottom) cells. Arrows indicate injection of metabolic modifiers to inhibit mitochondria (oligo/FCCP) and glycolysis (2-deoxyglucose: 2-DG). (C, D) Energy map showing baseline (<u>B</u>: black outlined) and maximal (<u>U</u>: red outlined) OCR and ECAR following OBM addition as fold change over uninduced ST2 (C) or MC3T3-E1 (D) cells. The four quadrants represent cell metabolism attributable to “Quiescent,” “Aerobic,” “Energetic,” or “Glycolytic” energy usage. Blue and orange colored ellipses represent the metabolic space covered by uninduced and induced cells, respectively. Inset shows the slope of each line generated from the xy-scatter plot. Slopes > 1 indicate metabolic shifts towards aerobic respiration, slopes <1 indicated metabolic shits towards glycolysis. (E) Specific metabolic contributions to OCR were calculated as Basal OCR (Baseline minus AA/Rot), ATP-production (Baseline minus Oligo), Respiratory capacity (FCCP minus baseline), Proton leak (Oligo minus AA/Rot), and Non-mitochondrial respiration (Post AA/Rot) represented as fold change from uninduced controls. (F) Specific metabolic contributions to ECAR calculated as Basal ECAR (Baseline minus 2-DG), Glycolytic Capacity (Oligo/FCCP minus Baseline), and Non-glycolytic acidification (Post 2-DG) represented as fold change from uninduced controls. Data are means ± SD (n=15). *, *p* <0.05 vs uninduced control as determined by ANOVA and confirmed by post hoc Bonferroni correction.

### Osteoinduction differentially stimulates OCR and ECAR in ST2 and MC3T3-E1 cells

The induction of osteogenic differentiation in ST2 and MC3T3-E1 cells resulted in a small but significant increase in mitochondrial engagement as measured by basal respiration without significantly increasing basal glycolytic rates (Figure 2E, F). ST2 cells rapidly induced doubling in both ATP-linked respiration and mitochondrial respiratory capacity (Figure 2E, F) and moderate, but significant increase in glycolytic capacity (Figure 2E, F). These data indicate a heightened flux of metabolic processes providing substrates for glycolysis and the mitochondrial electron transport chain in osteoinduced cells. Interestingly, the kinetics of osteoinduction in ST2 cells was markedly faster than those of MC3T3-E1 cells with the majority of the increase in OCR present after 1 hour of induction. MC3T3-E1 cells showed progressive increases in OCR parameters with the largest increase occurring after 12 hours. While ST2 cells demonstrated a more rapid response to OBM, MC3T3-E1 cells’ maximal response was larger. The proportion of OCR directly attributable to ATP production (i.e. Oligo sensitive OCR) was increased in proportion to increase in basal OCR despite an increase in proton leak (Oligo insensitive, AA/Rot sensitive OCR) and non-mitochondrial OCR (AA/Rot insensitive OCR) (Figure 2E, F). Taken together, these data demonstrate that these osteoprogenitor cells acutely respond to osteoinduction by increasing mitochondrial energetics. Additionally, these data suggests that osteoprogenitors from different lineages may have distinct metabolic profiles following osteoinduction.

### Biologically relevant osteoinducers, Wnt3a and BMP2, acutely activate mitochondrial bioenergetic parameters

Incubation of cells with ascorbic acid and β-glycerol phosphate is an accepted *in vitro* method of osteoinduction; however, it is not representative of physiologic induction. We therefore asked whether any cytokines or hormones known to promote osteogenic differentiation reproduced the acute response of OBM described in Figure 2. We observed that incubation of cells with IGF1, TGFβ, or PTH peptide (aa1-34) for 12 hours had no impact on the acute response of cells. In contrast, both BMP2 and Wnt3a strongly induced aerobic activation (Figure 3). Furthermore, this increase in OCR matched the profile we observed in OBM, i.e. mitochondrial respiratory flux, glycolytic reserve, and a proton leak all significantly increased (Figure 3B, C). Interestingly ATP-linked OCR did not change following BMP2 stimulation (Figure 3B). Additionally, we note that ST2 cells continue to display increased metabolic response over MC3T3-E1 cells as measured by their time to maximum response to osteoinduction, while MC3T3-E1 display a greater maximal increase. Using unbiased principle component analysis (PCA), we compared the changes in mitochondrial and glycolytic parameters over time and in response to OBM, BMP2, and Wnt3a to determine whether ST2 and MC3T3-E1 cells did indeed have distinct metabolic profiles following osteoinduction (Figure 4A). This analysis revealed partially overlapping yet diverging metabolic spaces occupied by each cell type. Unbiased heatmap clustering of these metabolic rates confirmed that activation of OCR is a common response to osteoinduction as the 12 hour post-induction data sets cluster together and have the greatest increase in OCR (Figure 4B). Additionally, cell-specific (ST2 and MC3T3-E1) responses showed the next greatest impact on cluster formation, with the method of osteoinduction having the least impact on how the data separated.

**Figure 3:**
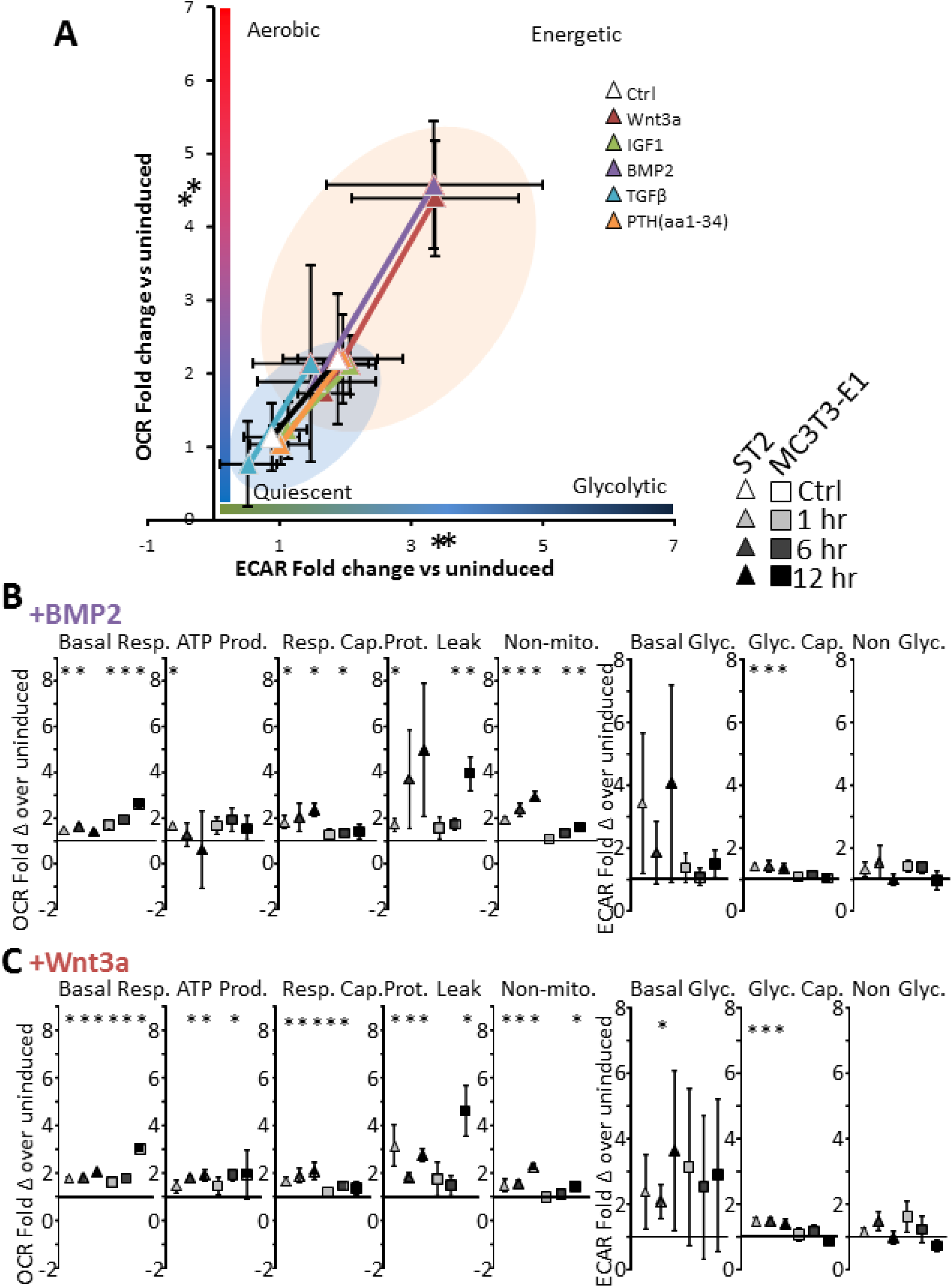
Acute activation of mitochondrial parameters is differently impacted by BMP2 and Wnt3a. (A) Energy map showing baseline (black outlined) and maximal (red outlined) OCR and ECAR as fold change over uninduced ST2 following addition of OB stimuli Wnt3a (red), IGF1 (green), BMP2 (purple), TGFβ (blue), or PTH (aa1-34) (orange). The four quadrants represent cell metabolism attributable to “Quiescent,” “Aerobic,” “Energetic,” or “Glycolytic” energy usage. Blue and orange colored ellipses represent the metabolic space covered by uninduced and induced cells, respectively. Specific metabolic changes caused by BMP2 (B) or Wnt3a (C) calculated as in Figure 3 and represented as fold change from uninduced controls. Data are means ± SD (n=15). *, *p* <0.05 vs uninduced control as determined by ANOVA and confirmed by post hoc Bonferroni correction.

**Figure 4:**
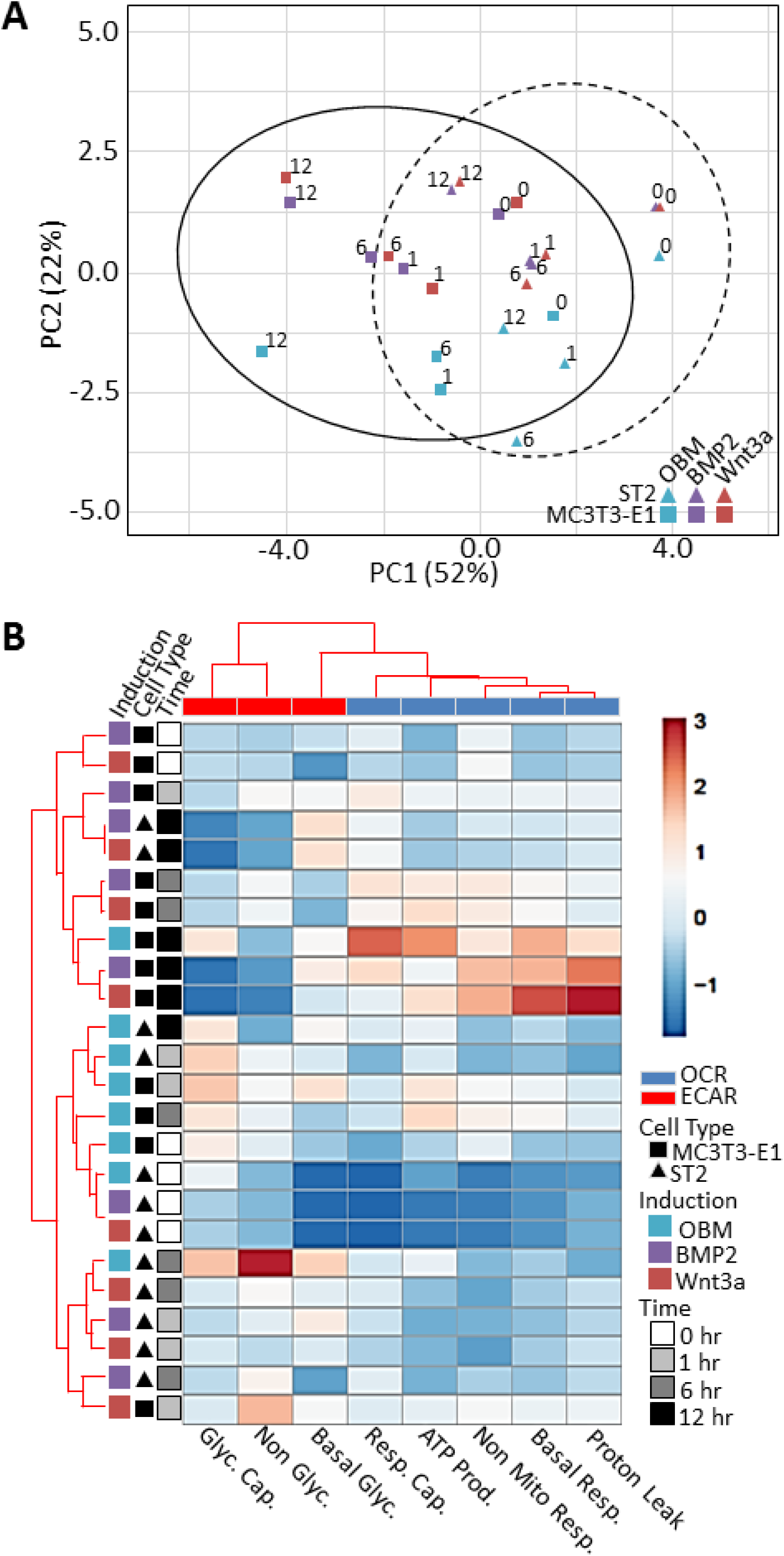
Unbiased comparison of cell type response to osteoinduction methods using key parameters of glycolytic and mitochondrial function as endpoints. (A) Principal component analysis (PCA) of metabolic parameters calculated from OCR and ECAR raw data for ST2 and MC3T3-E1 cells following osteoinduction with OBM, BMP2, and Wnt3a, at all four time points (0, 1, 6, 12 hours). The first two components show the data from the angle of most variability with PCA1 being cell type and PCA2 being osteoinducing factor. (B) Heatmap to visualize sample relations as a matrix. Values in the matrix are color coded to represent degree of difference between groups with dark red being largest increase and dark blue being largest decrease.

### Activation of Mitochondria requires Akt signaling

Mitochondrial activation can be the result of multiple signals and pathways, including bulk mitochondrial biogenesis, changes in cellular ratios of ATP/ADP, NAD^+^/NADH, calcium, metabolic intermediates of glycolysis and the Krebs’ cycle, and mitochondrial fusion/fission dynamics. The observed acute time scales of mitochondrial activation suggest that the response is mediated via changes in signaling pathways. One such signaling pathway implicated in both osteogenic differentiation (25), and regulation of mitochondrial function (26) is Akt. We therefore asked whether we could detect differences in Akt phosphorylation status following acute osteoinduction. Western blotting revealed increased Akt phosphorylation and thus activation in less than 1 hour using OBM, BMP2, or Wnt3a to induce the osteogenic program in ST2 and MC3T3-E1 cells (Figure 5A). Akt is known to have multiple phosphorylation targets in mitochondria (27)To check if Akt phosphorylated mitochondrial targets, we isolated mitochondria and performed western blotting of mitochondrial protein extracts from ST2 or MC3T3-E1 cells using anti-Akt phosphorylation sequence (RXXSp/Tp) antibody. The assay showed that mitochondria from both cell types contained increased levels of Akt-dependent phosphorylation following osteoinduction compared to mitochondria from uninduced cells (Figure 5B). MC3-T3 cells showed an increase in the pospohorylation of a protein approximates 50 kDa, whereas numerous bands were observed to increase in ST2 cells at molecular weights of approximately 100, 80, 60, 55, 50 and 30 kDa. It is work noting that Akt-dependent phosphorylation of mitochondrial proteins of the same molecular weights was described previously in mouse isogenic hepatocytes (28). While BMP2 appeared to have the strongest impact on increased band intensity, all three methods of osteoinduction showed increases over control. Densitometry histograms for each lane are shown to the right of each anti-phospho-Akt substrate blot. Such Akt-mediated phosphorylation of various mitochondrial targets is well documented to stimulate mitochondrial OxPhos activity [(27,29)]. These data indicate that all three methods of osteoinduction converge on Akt at the signaling level leading to Akt stimulatory effect on mitochondria. We next asked whether an Akt inhibitor could block the increased mitochondrial OCR observed in cells incubated in OBM. Osteoinduction of cells in the presence of a pan-Akt inhibitor completely and persistently blocked mitochondrial activation in both ST2 and MC3T3-E1 cells following osteoinduction so that there was no significant change in any mitochondrial bioenergetic parameter (Figure 6A). Importantly incubation of cells with a pan Akt inhibitor completely abrogated Akt activation, as evidenced by decreased phosphorylated-Akt signal (Figure 6B, C). This demonstrates that osteoinduction acts through Akt to promote mitochondrial bioenergetics. Finally, Akt inhibition prevented ST2 and MC3T3-E1 differentiation by OBM, and Wnt3a as evidenced by decreased ALP staining. Akt inhibition also prevented differentiation of ST2 cells by BMP2 and partially decreased MC3T3-E1 response to BMP2 (Figure 6D, E). Taken together our data demonstrate a model in which osteoinduction both via standard ascorbate/BGP treatment and physiologic signaling through BMP2 or Wnt3a results in activation of Akt, and increased Akt-mediated phosphorylation of mitochondrial proteins. Concurrently osteoinduced cells exhibit an Akt-dependent increase in mitochondrial bioenergetic parameters ultimately facilitating differentiation along the osteoblast lineage (Figure 7).

**Figure 5:**
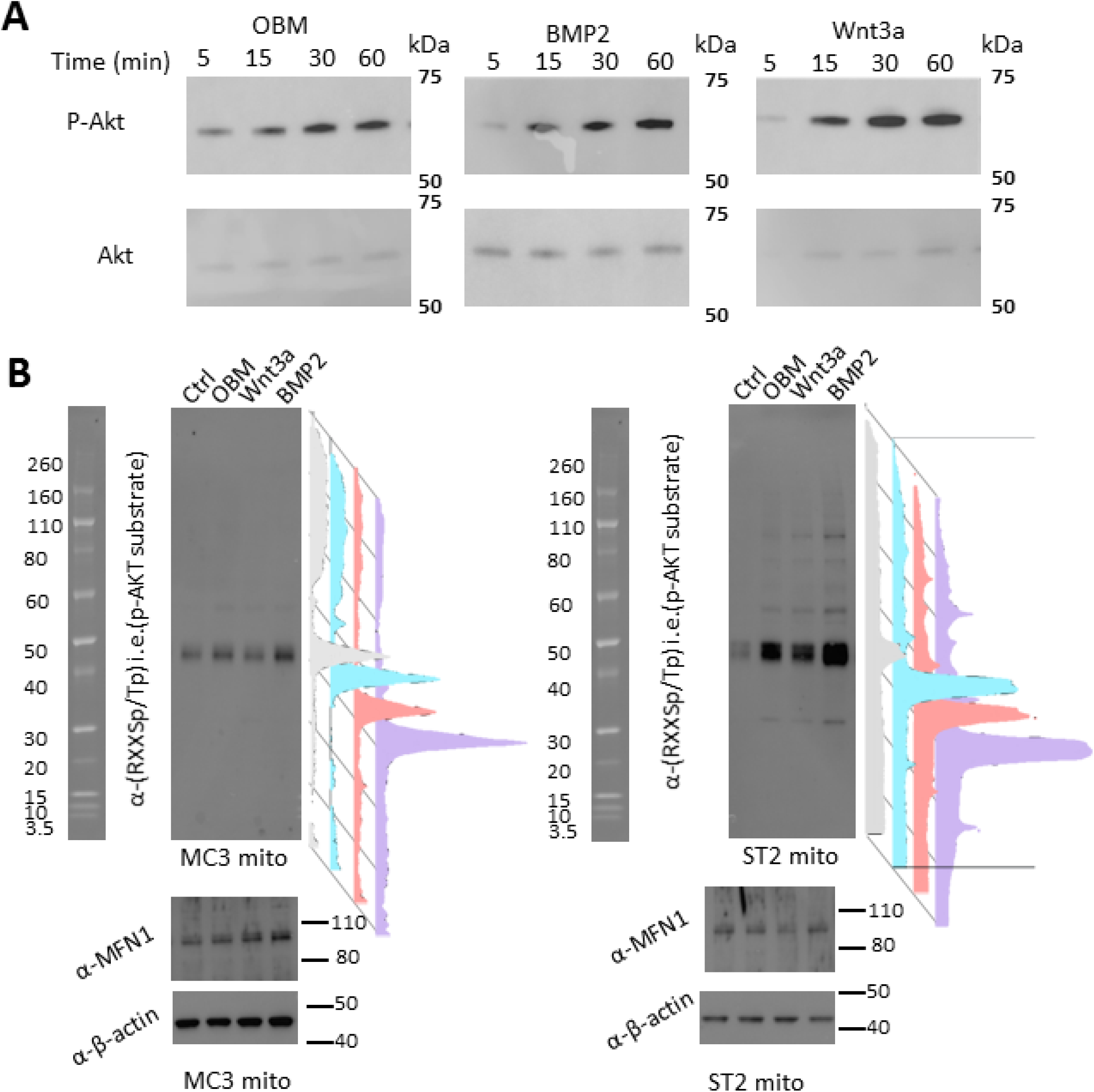
Osteoinduction activates Akt and increases phosphorylation of mitochondrial proteins. (A) Western blots of phospho-Akt and total Akt protein following 5, 15, 30 or 60 minutes incubation with OBM (left), BMP2 (center), or Wnt3a (right). (B) Western blot using anti-Akt phosphorylation sequence (RXXSp/Tp) to detect Akt target proteins in mitochondrial protein extracts following 48 hours osteoinduction, with mitochondrial loading control MFN1 and total protein loading control β-actin. Densitometry histograms showing relative intensity of bands from control (grey), OBM, (cyan), Wnt3a (rose), or BMP2 (lavendar). Blots are representative images of 3 replicates

**Figure 6:**
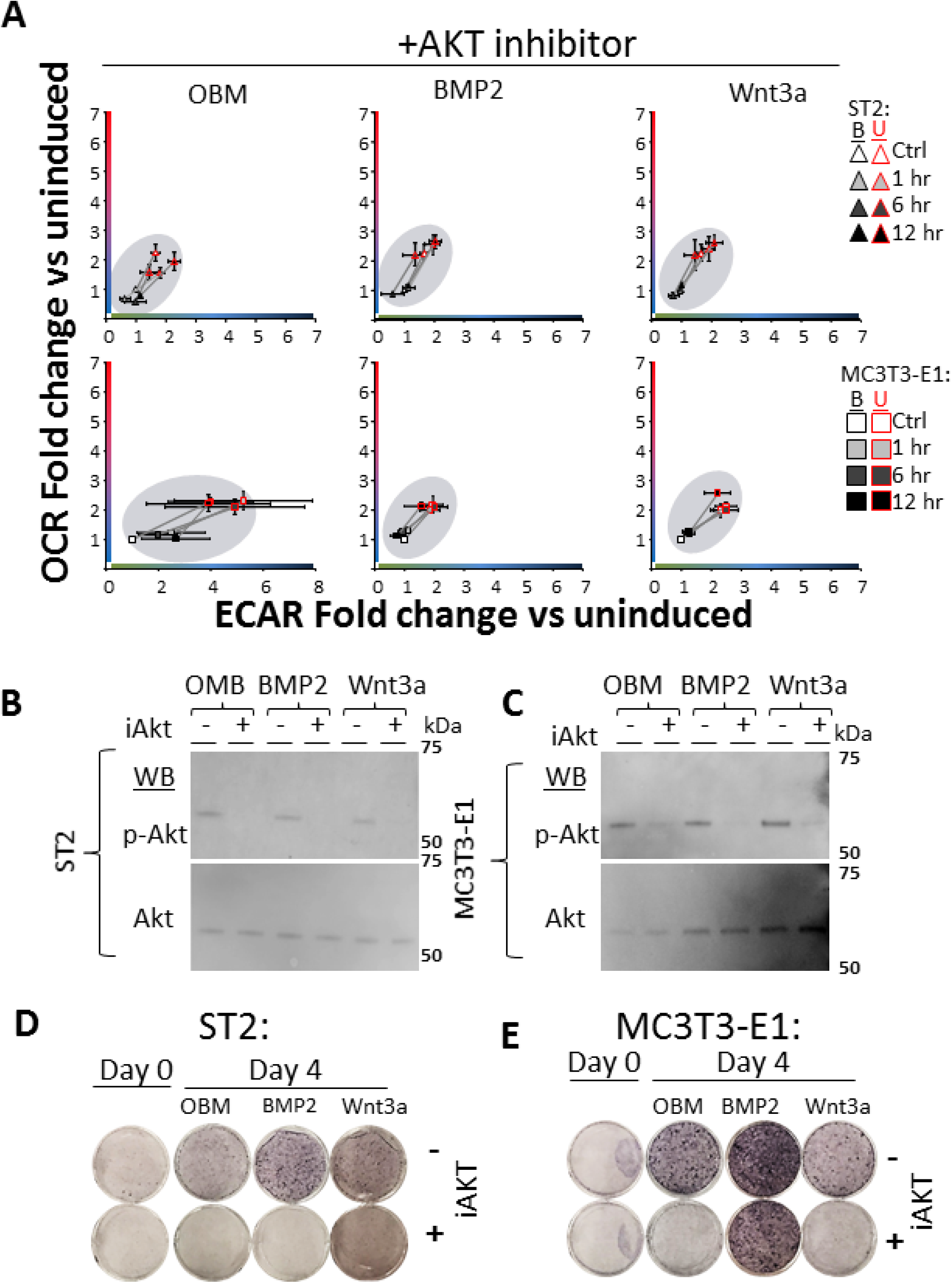
Activation of mitochondrial Spare capacity by osteo-induction requires active AKT. (A) Energy map of ST2 (top) and MC3T3-E1 (bottom) showing baseline (<u>B</u>: black outlined) and maximal (<u>U</u>: red outlined) OCR and ECAR of cells incubated with Akt inhibitor (iAkt) for 24 hours then induced using OBM (left), BMP2 (center), or Wnt3a (right) addition. Data represented as fold change over uninduced control cells. (B, C) Western blots of phospho-Akt and total Akt protein following incubation with iAkt for 24 hours then OBM (left), BMP2 (center), or Wnt3a (right) addition for 12 hours for ST2 (B) and MC3T3-E1 (C) cells. (D, E) ALP stain for cells treated with or without iAkt for 24 hours prior to OB induction with OBM, BMP2, or Wnt3a for four days. Seahorse n=15, significance by ANOVA and confirmed by post hot Bonferoni correction. (*) indicate p-value <0.05 vs uninduced control. Western blots and stains are representative images from n=3 replicates.

**Figure 7:**
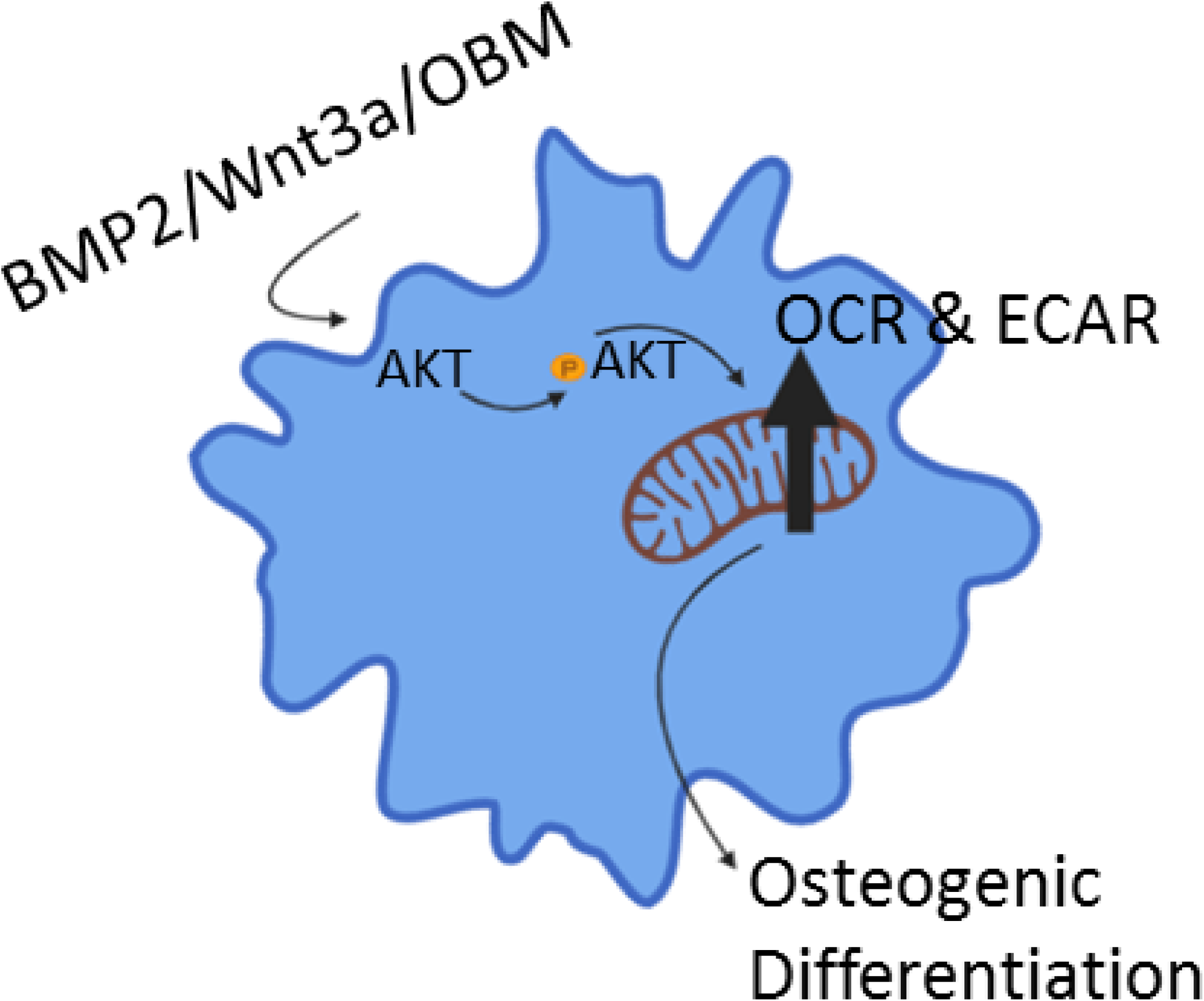
Schematic for Akt-dependent activation of mitochondrial function to induce osteogenic program. Standard acorbate/BGP treatment and physiologic signaling through BMP2 or Wnt3a results in activation of Akt, and increased Akt-mediated phosphorylation of mitochondrial proteins. Akt-dependent increase in mitochondrial bioenergetic parameters allows for differentiation along the osteoblast lineage. Figure generated using Biorender software: Biorender.com

## Discussion

The role of cellular metabolism during osteogenic differentiation is a topic of debate in the field. While it is routinely reported that various somatic stem cells rely on glycolysis for energy prior to differentiation and on the mitochondrial process of OxPhos during and after differentiation (4), the data regarding metabolism of osteoprogenitors remains controversial. Methodologic differences including supraphysiological levels of glucose and glutamine used in culture media to stimulate BMSC expansion have the potential to influence cell metabolism. The presence of extracellular pyruvate in metabolic assays could bypass any constraints that specifically affect OxPhos and artificially increase oxygen consumption in undifferentiated cells impeding the detection of change in OxPhos during differentiation. Furthermore, the presence of ascorbate in conventional media could prime osteogenic programming for activation and account for the alternate outcomes observed in reports which demonstrate increased OCR (10,11), and those that did not see changes on OCR (12,15,17).

In the current study, our experimental model used cells grown in minimal media lacking ascorbate and containing physiologic (1 g/L = 5 mM) glucose and heat inactivated FBS to best approximate physiologic conditions and to maintain cells in a true uninduced state. OCR and ECAR measures were performed in the absence of added pyruvate and with physiologic glucose and glutamine (1 mM). Additionally, the inclusion of the glutaminase inhibitor BPTES did not impact OCR or ECAR levels in a separate experiment (data not shown), indicating that glutamine is not a significant fuel source in pre-OB cells in this acute time frame following osteoinduction. Increased mitochondrial function can be a result of changes in the dynamic network of mitochondria within cells, changes in mitochondrial coupling, or activation of mitochondrial biogenesis. We chose time points on a short enough time scale to allow us to focus on changes in existent mitochondrial function rather than biogenesis of new mitochondrial protein machinery (30–32). Using these techniques, we observed rapid and consistent increases in basal mitochondrial OCR, mitochondrial directed ECAR (i.e. aerobic glycolysis), and respiratory capacity in both ST2 and MC3T3-E1 cells following addition of OBM, BMP2, or Wnt3a, as well as a consistent increase in proton leak. The method of osteoinduction resulted in distinct changes in different metabolic parameters with OBM and Wnt3a triggering an increase in ATP production while BMP2 did not. Furthermore, ST2 cells responded to osteoinduction early, with much of the maximal increase occurring in the first hour post-induction. In contrast MC3T3-E1 cells responded less rapidly, with the largest increase in metabolic parameters occurring at the 12-hour time points (compare Figures 3&4). Finally, ST2 cells overall fold increase in metabolic parameters while rapid, was smaller than the final increases observed in MC3T3-E1 cells. Taken together, ST2 and MC3T3-E1 cells appear to occupy distinct metabolic spaces both pre- and post-osteoinduction despite responding similarly to induction. Additionally, BMP2 and Wnt3a trigger metabolic activation of these cells with subtle differences that overlap with the response to OBM. This activation of mitochondrial parameters early in osteoinduction is essential and these cells are unable to tolerate even mild inhibition of mitochondrial electron transport chain complexes or ATP-synthase during this transition (Figure 1C, D). Interestingly mild uncoupling using FCCP did not negatively impact these cells. Mild uncoupling has been investigated as a therapeutic strategy in improving metabolic health and could be acting by a similar mechanism in osteoinduction as well (33).

A number of signaling pathways including Wnt3a and BMP2 are known to regulate osteoinduction of BMSCs into OBs (34). Our data demonstrate that Wnt3a and BMP2 can trigger an acute activation of mitochondrial OCR; and that interventions that block mitochondrial activation, prevent osteoinduction (Figure 1, 6). This agrees with the understanding of Wnt3a/β-catenin and BMP signaling as early signaling mechanisms during stem cell commitment stage (35). Additionally, post-translational modification of β-catenin modulates its activity. Active Wnt signaling initiates hypo-phosphorylation of β-catenin and decreased ubiquitination resulting in stabilization of the Wnt protein and ultimately the activation of expression of the canonical Wnt target genes. Phosphorylation promotes β-catenin degradation and is in competition with acetylation for amino acids in Wnt, which impacts the ubiquitination status of the protein. (36). Acetyl CoA (Ac-CoA) is a mitochondrially generated substrate for protein acetylation reactions by acetyltransferases. Thus, mitochondrially derived Ac-CoA may provide a link between cellular bioenergetic and differentiation pathways. Additionally acetylation has been shown to stabilize and promote activities of other key osteogenic factors, Runx2 and Osterix (37,38).

Osteoinduction has been previously reported to involve both a Wnt/Akt, and Bmp/Akt signaling axis, however these reports did not investigate mitochondrial parameters (39,40). Additionally, Akt has been shown separately to be an stimulator of mitochondrial function and an upstream regulator of the formation of mitochondria-associated membranes (26,29). Our data indicate Akt signaling as absolutely essential for the acute stimulation of mitochondrial OCR during the process of Wnt- and BMP-mediated osteoinduction.

It is interesting to note that while both ST2 and MC3T3-E1 cells had similar responses in our hands, their specific time to maximal response and the overall degree of mitochondrial activation were different. ST2 cells showed a rapid and large activation while MC3T3-E1 cells demonstrated a slower and more modest activation. As these cell lines are derived from osteoprogenitor cells of different embryonic origins, this difference in metabolic activity is not surprising and has also been observed in previous studies (14).

In conclusion, we found that: i) activation of mitochondrial OxPhos is achieved rapidly following induction of the osteogenic program in both long bone- and calvarial bone-derived osteoprogenitor cells; (ii) mitochondrial respiratory activity is required for successful transition into the osteogenic program; and (iii) activation of mitochondria using these methods is dependent upon active Akt signaling (Figure 7). We, therefore suggest the scenario wherein the osteoprogenitor mitochondria are rapidly activated following Akt phosphorylation of mitochondrial targets allowing for the subsequent activation of the osteogenic program.

## Materials and Methods

### Cell Culture

Mouse long bone-derived ST2 cells were a gift from Dr. Clifford Rosen. Mouse calvarial bone-derived MC3T3-E1 cells were acquired from ATCC. Cells were expanded in sterile conditions and maintained in a 37°C incubator at 5% CO_2_ in αMEM media (Gibco A10490-01) containing physiologic 1 g/L (i.e. 5 mM) glucose, 0.3 g/L (i.e. 1 mM) L-glutamine, ribonuclesides (0.01 g/L each), deoxyribonucleosides (0.01 g/L each), no ascorbic acid, 10% FBS (Gibco 10437-028) heat inactivated for 30 minutes at 55 degrees C, and 0.1 % Penn/Strep (Gibco 15140-122). Cells were maintained at passage numbers < 20 as recommended by the supplier and from previous handling experience.

### Osteoinduction and detection by cell staining

The addition of 50 μg/mL ascorbate (Sigma A4544) and 2.5 mM β-glycerol phosphate (USB Corp Cleveland, OH, 21655), 25 ng/mL BMP2 (R&D systems 355-BM-050/CF), 25 ng/mL Wnt3a (R&D systems 5036-WN-010), 5 ng/mL IGF1 (Sigma 13769-50UG), 0.2 ng/mL TGFβ (R&D systems 240-B), or 1 ng/mL PTH aa1-34 (R&D systems 3011/1) to αMEM media induced osteogenesis. To confirm osteogenesis, cells were stained with OB-specific ALP-specific stain (Thermo NBT/BCIP 1-step 34042) and with 0.5 % CV (Sigma C3886) to determine total cell count as previously described (13). For metabolic inhibitory studies, cells were incubated with either 0.1 μg/mL Oligomycin (Oligo, Sigma 75351), 0.1 μM Antimycin A (AA, Sigma A8674), 0.1 μM Rotenone (ROT, Sigma R-8875), or 0.2 μM Carbonyl cyanide 4-(trifluoromethoxy)phenylhydrazone (FCCP, Sigma C2920) for 48 hours prior to osteoinduction, and inhibitors were present during the entire time course of induction. For Akt signaling studies, cells were incubated with 10 μM Akt1/2 inhibitor (Sigma A6730) for 24 hours prior to osteoinduction (or as indicated for time course studies) and were present during the period of osteoinduction.

### Metabolic profiling

OCR and ECAR were measured using Seahorse XF96 (Seahorse Bioscience). Cells were plated on Seahorse 96-well plates 48 hours before the experiment at a density of 15,000 cells/well. Cells were incubated in media containing control, OBM, BMP2 or Wnt3a to induce osteogenesis for 1, 6, or 12 hours. Immediately before the experiment, medium was replaced with unbuffered DMEM (Sigma D5030) medium containing 1 mM glutamine (Gibco 25030-081), 5 mM glucose (Sigma G8270), appropriate osteogenic inducer, and no pyruvate (pH 7.4). A baseline measurement of OCR and ECAR was taken, and then an inhibitory analysis was performed using serial injections of 1 μg/mL Oligo, 2 μM FCCP, and simultaneous addition of 1 μM each of AA and ROT. A parallel experiment was run with the following injections of Oligo+FCCP followed by 15 mM 2-deoxyglucose (Sigma D-3179). The following OxPhos and glycolytic indexes were calculated: basal respiration (OCR_pre-Oligo_ − OCR_post-AntA_), ATP-linked respiration (OCR_pre-Oligo_ − OCR_post-Oligo_), spare respiratory capacity (OCR_post-FCCP_ − OCR_pre-Oligo_), proton leak (OCR_post-Oligo_ − OCR_post-AA/ROT_), basal glycolysis (ECAR_pre-Oligo_), and glycolytic flux (ECAR_post-Oligo/FCCP_ – ECAR _pre-oligo/FCCP_).

### Real-time RT-PCR

Total RNA was isolated using the RNeasy kit (Qiagen 74106) and reverse transcribed into cDNA using the qScript cDNA synthesis kit (Quanta 95048-500). cDNA was subjected to real-time RT-PCR. The primer pairs used for genes of interest are outlined in Table 1. Real-time RT-PCR was performed in the RotorGene system (Qiagen) using SYBR Green (Quanta 95072-012). The expression of genes of interest was normalized to expression of beta-2 microglobulin (*B2m*).

**Table 1.**
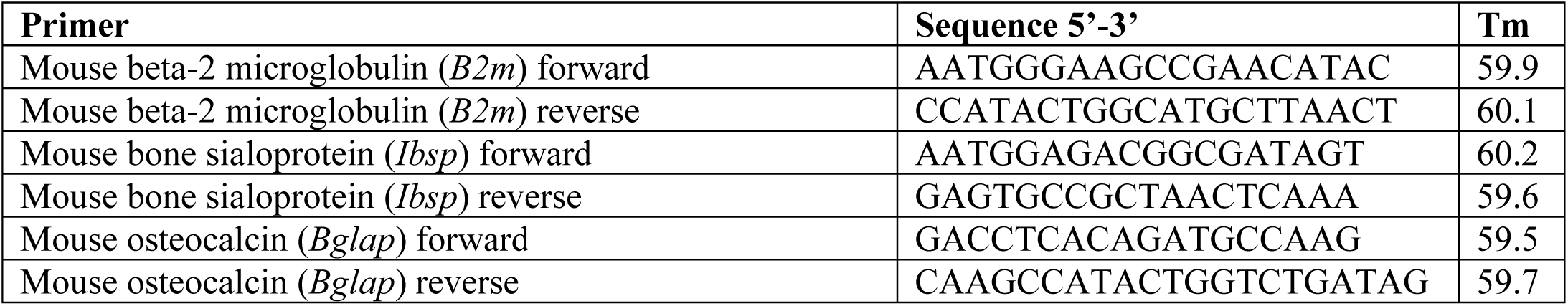

### Western blotting

Cells were lysed with lysis buffer containing protease inhibitors and subjected to 4–12% SDS-PAGE followed by transfer to polyvinylidene difluoride membranes (PVDF) and blocking in 5% dry milk. For Akt detection, blots were probed total Akt antibody (Cell Signaling 92728) at a concentration of 1:2000 For phosphorylated Akt detection, blots were probed with mouse monoclonal phospho serine 473 Akt (Cell Signaling 4060S) at a concentration of 1:1000. Akt substrates were detected using p-Akt Substrate RXXS*/T* (Cell Signaling 110B7E) at 1:2000. Loading controls were detected using antibodies against: 1:2000 Mitofusin 1 (Abcam Ab57602), 1:20,000 β-actin (Invitrogen Cat-15). Following wash of primary antibody, blots were incubated with either horseradish peroxidase (HRP)–conjugated goat anti-mouse antibody (Bio-Rad 170-6516) or HRP–conjugated goat anti-rabbit antibody at a concentration of 1:5000. Antibody detection was developed with West Femto Substrate (Thermo Scientific 34095). Densitometry was preformed using Fiji software “gel analysis” plug-in.

### Mitochondrial Isolation

Cells were grown on 15cm plates until confluency and treated for 48 hours with osteogenic media. Cells were rapidly transferred to cold PBS containing phosphatase inhibitors by scraping (all following steps contain phosphatase inhibitors). Cells were spun down at 3,500 rpm then resuspended in MannPrep buffer (Mannitol at 195 mM, Sucrose at 65 mM, HEPES at 2 mM, KH_2_PO_4_ at 0.05 mM, EGTA at 0.05 mM, MgCl_2_ at 0.01 mM; pH 7.4 at 4°C). Cells were broken up by homogenizing them in glass homogenizer with teflon pestle and 35 passes. Crude precipitate and nuclear material was removed by spin at 3,500 rpm for 5 min at 4°C in benchtop centrifuge. Supernatant was removed and spun at 12,000 rpm for 9 min at 4°C in benchtop centrifuge to pellet mitochondria. Crude mitochondrial pellet was resuspended in 0.02 ml protein lysis buffer.

### Statistical analysis

Mean values and standard deviation were calculated, and the statistical significance (*p* < 0.05) was established using either Student’s *t*-test when two variables were compared or one-way analysis of variance (ANOVA) with post hoc Bonferroni correction when more than two variables were compared based on normal spread of our data as indicated in the figure legends. Principle component analysis (PCA) and heat map were performed using open access ClustVis software (41,42).

## FOOTNOTES

Funding was provided by the National Institute of Health grant to R.A.E. (R01 AR072601).

## The abbreviations used are

AA: Antimycin A
ALP: Alkaline phosphatase
BMP2: Bone Morphogenic Protein 2
CV: Crystal Violate
ECAR: Extracellular Acidification Rate
FCCP: Carbonyl cyanide 4-(trifluoromethoxy)phenylhydrazone
OBM: Osteoblastic media a.k.a. Ascorbic acid and beta-gycero phosphate
OBs: Osteoblasts
OCR: Oxygen Consumption Rate
Oligo: Oligomycin
OxPhos: Oxidative Phosphorylation
ROS: Reactive Oxygen Species
ROT: Rotenone
Wnt3a: Wingless protein 3a

---

**Table 1.**
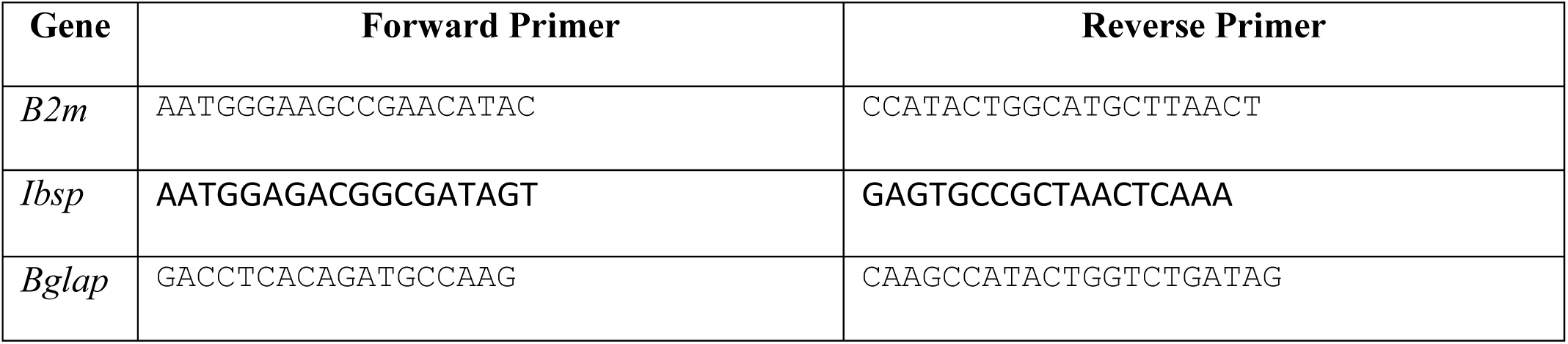
Primers used for real time RT-PCR analysis. All primers written from 5’ to 3’ (left to right).

## References

1. Nunnari J, Suomalainen A. Mitochondria: in sickness and in health. Cell [Internet]. 2012 Mar 16 [cited 2019 Oct 22];148(6):1145–59. Available from: http://www.ncbi.nlm.nih.gov/pubmed/22424226

2. McBride HM, Neuspiel M, Wasiak S. Mitochondria: More Than Just a Powerhouse. vol. 16, Current Biology. 2006.

3. Schorr S, van der Laan M. Integrative functions of the mitochondrial contact site and cristae organizing system. vol. 76, Seminars in Cell and Developmental Biology. Elsevier Ltd; 2018. p. 191–200.

4. Varum S, Rodrigues AS, Moura MB, Momcilovic O, Easley IV CA, Ramalho-Santos J, et al. Energy metabolism in human pluripotent stem cells and their differentiated counterparts. PLoS One. 2011;6(6).

5. Sandhir R, Halder A, Sunkaria A. Mitochondria as a centrally positioned hub in the innate immune response. vol. 1863, Biochimica et Biophysica Acta – Molecular Basis of Disease. Elsevier B.V.; 2017. p. 1090–7.

6. Cherry C, Thompson B, Saptarshi N, Wu J, Hoh J. 2016: A “Mitochondria” Odyssey. Trends Mol Med [Internet]. 2016 May [cited 2019 Oct 15];22(5):391–403. Available from: http://www.ncbi.nlm.nih.gov/pubmed/27151392

7. Vafai SB, Mootha VK. Mitochondrial disorders as windows into an ancient organelle. Nature [Internet]. 2012 Nov 15 [cited 2019 Oct 15];491(7424):374–83. Available from: http://www.ncbi.nlm.nih.gov/pubmed/23151580

8. Bianco P, Cao X, Frenette PS, Mao JJ, Robey PG, Simmons PJ, et al. The meaning, the sense and the significance: translating the science of mesenchymal stem cells into medicine. Nat Med [Internet]. 2013 Jan [cited 2019 Oct 15];19(1):35–42. Available from: http://www.ncbi.nlm.nih.gov/pubmed/23296015

9. Motyl KJ, Guntur AR, Carvalho AL, Rosen CJ. Energy Metabolism of Bone. vol. 45, Toxicologic Pathology. SAGE Publications Inc.; 2017. p. 887–93.

10. Chen C-T, Shih Y-R V, Kuo TK, Lee OK, Wei Y-H. Coordinated changes of mitochondrial biogenesis and antioxidant enzymes during osteogenic differentiation of human mesenchymal stem cells. Stem Cells [Internet]. 2008 Apr [cited 2019 Oct 15];26(4):960–8. Available from: http://www.ncbi.nlm.nih.gov/pubmed/18218821

11. Guntur AR, L. PT, Farber CR, Rosen CJ. Bioenergetics during calvarial osteoblast differentiation reflect strain differences in bone mass. Endocrinology. 2014;155(5):1589–95.

12. Shum LC, White NS, Mills BN, Bentley KL de M, Eliseev RA. Energy Metabolism in Mesenchymal Stem Cells During Osteogenic Differentiation. Stem Cells Dev [Internet]. 2016 Jan 15 [cited 2019 Oct 15];25(2):114–22. Available from: http://www.ncbi.nlm.nih.gov/pubmed/26487485

13. Shares BH, Busch M, White N, Shum L, Eliseev RA. Active mitochondria support osteogenic differentiation by stimulating-Catenin acetylation. J Biol Chem. 2018;293(41):16019–27.

14. Quarto N, Wan DC, Kwan MD, Panetta NJ, Li S, Longaker MT. Origin matters: Differences in embryonic tissue origin and Wnt signaling determine the osteogenic potential and healing capacity of frontal and parietal calvarial bones. J Bone Miner Res. 2010 Jul;25(7):1680–94.

15. Esen E, Chen J, Karner CM, Okunade AL, Patterson BW, Long F. WNT-LRP5 signaling induces warburg effect through mTORC2 activation during osteoblast differentiation. Cell Metab. 2013 May 7;17(5):745–55.

16. Guntur AR, Gerencser AA, L. PT, DeMambro VE, Bornstein SA, Mookerjee SA, et al. Osteoblast-like MC3T3-E1 Cells Prefer Glycolysis for ATP Production but Adipocyte-like 3T3-L1 Cells Prefer Oxidative Phosphorylation. J Bone Miner Res. 2018 Jun 1;33(6):1052–65.

17. Pattappa G, Heywood HK, de Bruijn JD, Lee DA. The metabolism of human mesenchymal stem cells during proliferation and differentiation. J Cell Physiol [Internet]. 2011 Oct [cited 2019 Oct 15];226(10):2562–70. Available from: http://www.ncbi.nlm.nih.gov/pubmed/21792913

18. Komarova S V, Ataullakhanov FI, Globus RK. Bioenergetics and mitochondrial transmembrane potential during differentiation of cultured osteoblasts. Am J Physiol Cell Physiol [Internet]. 2000 Oct [cited 2019 Oct 15];279(4):C1220–9. Available from: http://www.ncbi.nlm.nih.gov/pubmed/11003602

19. Yu Y, Newman H, Shen L, Sharma D, Hu G, Mirando AJ, et al. Glutamine Metabolism Regulates Proliferation and Lineage Allocation in Skeletal Stem Cells. Cell Metab. 2019 Apr 2;29(4):966–978.e4.

20. Ganz J, Kaslin J, Freudenreich D, Machate A, Geffarth M, Brand M. Subdivisions of the adult zebrafish subpallium by molecular marker analysis Julia Ganz. J Comp Neurol [Internet]. 2011;520(Sfb 655):633–55. Available from: http://www.ncbi.nlm.nih.gov/pubmed/21858823

21. Forni MF, Peloggia J, Trudeau K, Shirihai O, Kowaltowski AJ. Murine Mesenchymal Stem Cell Commitment to Differentiation Is Regulated by Mitochondrial Dynamics. Stem Cells [Internet]. 2016 Mar [cited 2019 Oct 15];34(3):743–55. Available from: http://www.ncbi.nlm.nih.gov/pubmed/26638184

22. Otsuka E, Yamaguchi A, Hirose S, Hagiwara H. Characterization of osteoblastic differentiation of stromal cell line ST2 that is induced by ascorbic acid. Am J Physiol Physiol [Internet]. 1999 Jul 1 [cited 2019 Oct 15];277(1):C132–8. Available from: https://www.physiology.org/doi/10.1152/ajpcell.1999.277.1.C132

23. Wang D, Christensen K, Chawla K, Xiao G, Krebsbach PH, Franceschi RT. Isolation and Characterization of MC3T3-E1 Vivo Differentiation / Mineralization Potential. J Bone Miner Res. 1999;14(6):893–903.

24. Rastogi A, Joshi P, Contreras E, Gama V. Remodeling of mitochondrial morphology and function: an emerging hallmark of cellular reprogramming. Cell Stress. 2019 Jun 10;3(6):181–94.

25. Raucci A, Bellosta P, Grassi R, Basilico C, Mansukhani A. Osteoblast proliferation or differentiation is regulated by relative strengths of opposing signaling pathways. J Cell Physiol [Internet]. 2008 May [cited 2019 Oct 15];215(2):442–51. Available from: http://www.ncbi.nlm.nih.gov/pubmed/17960591

26. Betz C, Stracka D, Prescianotto-Baschong C, Frieden M, Demaurex N, Hall MN. MTOR complex 2-Akt signaling at mitochondria-associated endoplasmic reticulum membranes (MAM) regulates mitochondrial physiology. Proc Natl Acad Sci U S A. 2013 Jul 30;110(31):12526–34.

27. Bijur GN, Jope RS. Rapid accumulation of Akt in mitochondria following phosphatidylinositol 3-kinase activation. J Neurochem. 2003;87(6):1427–35.

28. Li C, Li Y, He L, Agarwal AR, Zeng N, Cadenas E, et al. PI3K/AKT signaling regulates bioenergetics in immortalized hepatocytes. Free Radic Biol Med. 2013 Jul;60:29–40.

29. Chen YH, Su CC, Deng W, Lock LF, Donovan PJ, Kayala MA, et al. Mitochondrial Akt Signaling Modulated Reprogramming of Somatic Cells. Sci Rep. 2019 Dec 1;9(1):1–14.

30. Scarpulla RC. Transcriptional paradigms in mammalian mitochondrial biogenesis and function. vol. 88, Physiological Reviews. 2008. p. 611–38.

31. Gottlieb RA, Bernstein D. Mitochondrial remodeling: Rearranging, recycling, and reprogramming. Cell Calcium [Internet]. 2016 Aug [cited 2017 Jan 5];60(2):88–101. Available from: http://linkinghub.elsevier.com/retrieve/pii/S0143416016300495

32. Wilson-Fritch L, Burkart A, Bell G, Mendelson K, Leszyk J, Nicoloro S, et al. Mitochondrial Biogenesis and Remodeling during Adipogenesis and in Response to the Insulin Sensitizer Rosiglitazone. Mol Cell Biol. 2003 Feb 1;23(3):1085–94.

33. M. Cunha F, C. Caldeira da Silva C, M. Cerqueira F, J. Kowaltowski A. Mild Mitochondrial Uncoupling as a Therapeutic Strategy. Curr Drug Targets. 2011 May 2;12(6):783–9.

34. Marie PJ. Transcription factors controlling osteoblastogenesis. Arch Biochem Biophys [Internet]. 2008 May 15 [cited 2019 Oct 15];473(2):98–105. Available from: http://www.ncbi.nlm.nih.gov/pubmed/18331818

35. Valenti MT, Dalle Carbonare L, Mottes M. Osteogenic Differentiation in Healthy and Pathological Conditions. Int J Mol Sci [Internet]. 2016 Dec 27 [cited 2019 Oct 15];18(1). Available from: http://www.ncbi.nlm.nih.gov/pubmed/28035992

36. Gao C, Xiao G, Hu J. Regulation of Wnt/β-catenin signaling by posttranslational modifications. Cell Biosci [Internet]. 2014 Mar 4 [cited 2019 Oct 15];4(1):13. Available from: http://www.ncbi.nlm.nih.gov/pubmed/24594309

37. Jeon EJ, Lee KY, Choi NS, Lee MH, Kim HN, Jin YH, et al. Bone morphogenetic protein-2 stimulates Runx2 acetylation. J Biol Chem. 2006 Jun 16;281(24):16502–11.

38. Lu J, Qu S, Yao B, Xu Y, Jin Y, Shi K, et al. Osterix acetylation at K307 and K312 enhances its transcriptional activity and is required for osteoblast differentiation. Oncotarget [Internet]. 2016 Jun 21 [cited 2019 Oct 15];7(25):37471–86. Available from: http://www.ncbi.nlm.nih.gov/pubmed/27250035

39. Eiraku N, Chiba N, Nakamura T, Amir MS, Seong C, Ohnishi T, et al. BMP9 directly induces rapid GSK3-β phosphorylation in a Wnt-independent manner through class I PI3K-Akt axis in osteoblasts. FASEB J [Internet]. 2019 Nov 1 [cited 2020 Jun 8];33(11):12124–34. Available from: https://onlinelibrary.wiley.com/doi/abs/10.1096/fj.201900733RR

40. Ling L, Dombrowski C, Foong KM, Haupt LM, Stein GS, Nurcombe V, et al. Synergism between Wnt3a and heparin enhances osteogenesis via a phosphoinositide 3-kinase/Akt/RUNX2 pathway. J Biol Chem. 2010 Aug 20;285(34):26233–44.

41. Metsalu T, Vilo J. ClustVis: A web tool for visualizing clustering of multivariate data using Principal Component Analysis and heatmap. Nucleic Acids Res. 2015;43(W1):W566–70.

42. Jolliffe I. Principal Component Analysis. In: Wiley StatsRef: Statistics Reference Online [Internet]. Chichester, UK: John Wiley & Sons, Ltd; 2014 [cited 2019 Oct 24]. Available from: http://doi.wiley.com/10.1002/9781118445112.stat06472

